# A planar neuromuscular controller to simulate compensation strategies in the sit-to-walk movement

**DOI:** 10.1101/2023.11.24.568552

**Authors:** Eline van der Kruk, Thomas Geijtenbeek

## Abstract

Standing up from a chair is a key daily life activity that is sensitive to functional limitations as we age and associated with falls, frailty, and institutional living. Predictive neuromusculoskeletal models can potentially shed light on the interconnectivity and interdependency of age-related changes in neuromuscular capacity, reinforcement schemes, sensory integration, and adaptation strategies during stand-up. Most stand-up movements transfer directly into walking (sit-to-walk). The aim of this study was to develop and validate a neuromusculoskeletal model with reflex-based muscle control that enables simulation of the sit-to-walk movement under various conditions (seat height, foot placement). We developed a planar sit-to-walk musculoskeletal model (11 degrees-of-freedom, 20 muscles) and neuromuscular controller, consisting of a two-phase stand-up controller and a reflex-based gait controller. The stand-up controller contains generic neural pathways of delayed proprioceptive feedback from muscle length, force, velocity, and upper-body orientation (vestibular feedback) and includes both monosynaptic an antagonistic feedback pathways. The control parameters where optimized using a shooting-based optimization method, based on a high-level optimization criterium. Simulations were compared to recorded kinematics, ground reaction forces, and muscle activation. The simulated kinematics resemble the measured kinematics and muscle activations. The adaptation strategies that resulted from alterations in seat height, are comparable to those observed in adults. The simulation framework and model are publicly available and allow to study age-related compensation strategies, including reduced muscular capacity, reduced neural capacity, external perturbations, and altered movement objectives.

## 1. Introduction

Adults rise from a seated position approximately 60 times a day (1). The inability to perform this task (e.g. from a seat or toilet) is associated with increased number of falls, frailty and need for institutional living(2). As the human movement system has physiological and functional redundancy, movement limitations do not promptly arise at the onset of physical decline (3,4). This makes these limitations difficult to anticipate, as it is unclear how much decline can be tolerated before movement limitations arise.

Humans adapt their movement strategies throughout the lifespan. They develop compensatory approaches to manage declines in neuromusculoskeletal capacity, changes in reinforcement schemes (such as prioritizing pain avoidance or stability due to fear of falling), and alterations in sensory input (Fig. 1) (5). In a recent study (6), we observed that increased neural delays are related to how humans adapt their standing up strategy as they age. However, confirming this solely through experimental studies is challenging due to the interconnectivity with other factors affecting neuromuscular capacity, such as reduced muscular strength or contraction velocity Neuromusculoskeletal simulations have the potential to shed light on the interconnectivity and interdependency of neuromuscular capacity and adaptation during stand up. Unlike in experimental studies, they allow researchers to isolate specific variables of age-related decline. However, we need a model and controller that can realistically simulate compensation strategies in the standing up movement before we can introduce reduced neuromuscular capacity and altered movement objectives.

**Figure 1:**
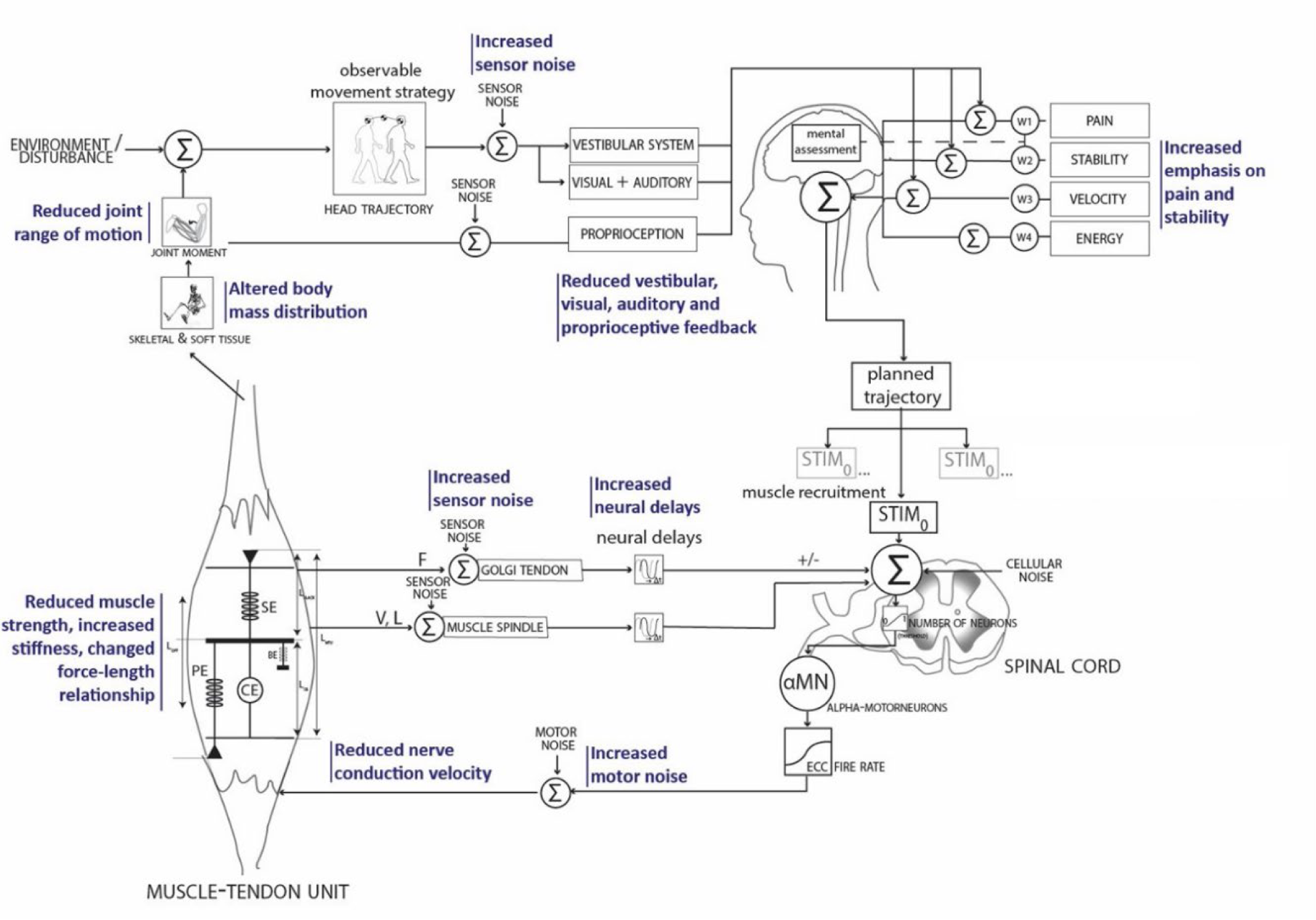
Feedback control of the neuromuscular system. In blue the parts of the neuromuscular system that start are known to have age-related changes (19).

Data driven simulations rely on experimental data, like the studies of Caruthers et al. (2016) (7) and Smith et al. (2020) (8) who used an inverse musculoskeletal approach to examine the individual muscle contributions to the sit-to-stand transfer. However, to study compensation strategies, we require simulations capable of generating movements “de novo” without measured inputs, which is achieved in predictive simulations. Predictive simulations can be broadly classified into two approaches: those that determine (open-loop) muscle controls, often known as trajectory optimization, such as direct collocation, and those that determine control policies, such as reflex-based controllers (9).

Trajectory optimization, such as the study of Bobbert et al. (2016) (10) on the sit-to-stand movement, is not suitable for simulation of neuromuscular adaptation strategies. These methods lack a neuromuscular feedback control model, which is required to simulate the effect of sensorimotor noise, external perturbations, or increased neural delays. Sensorial feedback control models solve for control policy and are a promising approach to simulate human movement based on the principles of the neural control system. Geyer and Herr (2010) (11) were pioneers to develop a robust gait controller with a limited set of proprioceptive feedback loops, a method subsequently applied in various gait simulation studies (12–14). These reflex controllers have been sporadically studied beyond gait, with Muñoz et al. (2022) using the controller to simulate a sit-to-stand motion (15). This existing sit-to-stand feedback controller has several limitations when it comes to realistic simulation of adaptation strategies in standing up(15). Asymmetry in muscular activation and kinematics is common in STS, especially in an elderly or patient population (7,16,17), but was neglected. Concerning the neural controller, it incorporates proprioceptive muscle length feedback via monosynaptic paths and vestibular feedback, but it neglects potential antagonistic paths and proprioceptive feedback related to force and velocity, crucial elements of neural organization. Additionally, the transitions between controller states are based on seemingly arbitrary kinematic thresholds.

The objective of our study was to develop and validate a neuromusculoskeletal model based on a generic reflex-based muscle control system, allowing simulation of the sit-to-walk movement under different conditions, such as seat height and foot placement. To our knowledge, this is the first predictive controller for simulating the sit-to-walk movement, which is a crucial aspect of the timed-up-and-go test used in clinical assessments. Our aim was to design a framework capable of simulating the effects of age-related decline in muscular and neural capacities, changes in movement objectives, and responses to external perturbations, as depicted in Figure 1, and therefore we specifically chose a control policy approach. The stand-up controller uses delayed proprioceptive feedback from muscle length, force, velocity, and upper-body orientation (vestibular feedback) and includes monosynaptic an antagonistic feedback pathways. The movement consists of two phases, each of which uses a separate set of feedback gains, as well as a separate set of excitation offsets. The transitions between the states are not prescribed but included as optimization parameter. The simulations were compared to recorded kinematics, ground reaction forces (GRF), and muscle activation (sEMG). The simulation framework and model are freely available (18).

## 2. Materials & Method

Our sit-to-walk controller consists of a two-phase stand-up controller (*P1, P2*) and a gait controller (*GAIT*) based on (11). The two-phase stand-up controller is based on proprioceptive feedback from muscle length, velocity, and force feedback, vestibular feedback and constant excitation (*C*), and uses separate feedback and excitation parameters for each phase. The feedback contains neural delays and includes both monosynaptic an antagonistic pathways. The free parameters were optimized to minimize the cubed muscle activation and gross cost of transport, at a prescribed minimum gait velocity, while avoiding falling, and excessive ligament stress. We ran multiple parallel optimizations with the same initial guess and used the best results as a starting point for the next round of optimizations. Final results were compared to a subset of recorded kinematics, ground reaction forces (GRF), and muscle activation (sEMG), in which participants were asked to stand up and walk to a table at self-selected speed (6). The simulations and optimizations were performed in SCONE (23) using the Hyfydy simulation engine (20).

### 2.1 Musculoskeletal model (H1120)

The musculoskeletal model (*H1120*) represents a male adult with a height of 1.80m and a mass of 75 kg optimized for predictive simulations (Fig. 2). The model has 11 degrees of freedom (dof): 3 dofs between the pelvis and the ground, a pin joint (1-dof) at the hip, ankle and knee, a 1-dof lumbar joint, between the pelvis and the lumbar, and a 1-dof thoracic joint between the lumbar and the torso.

**Figure 2:**
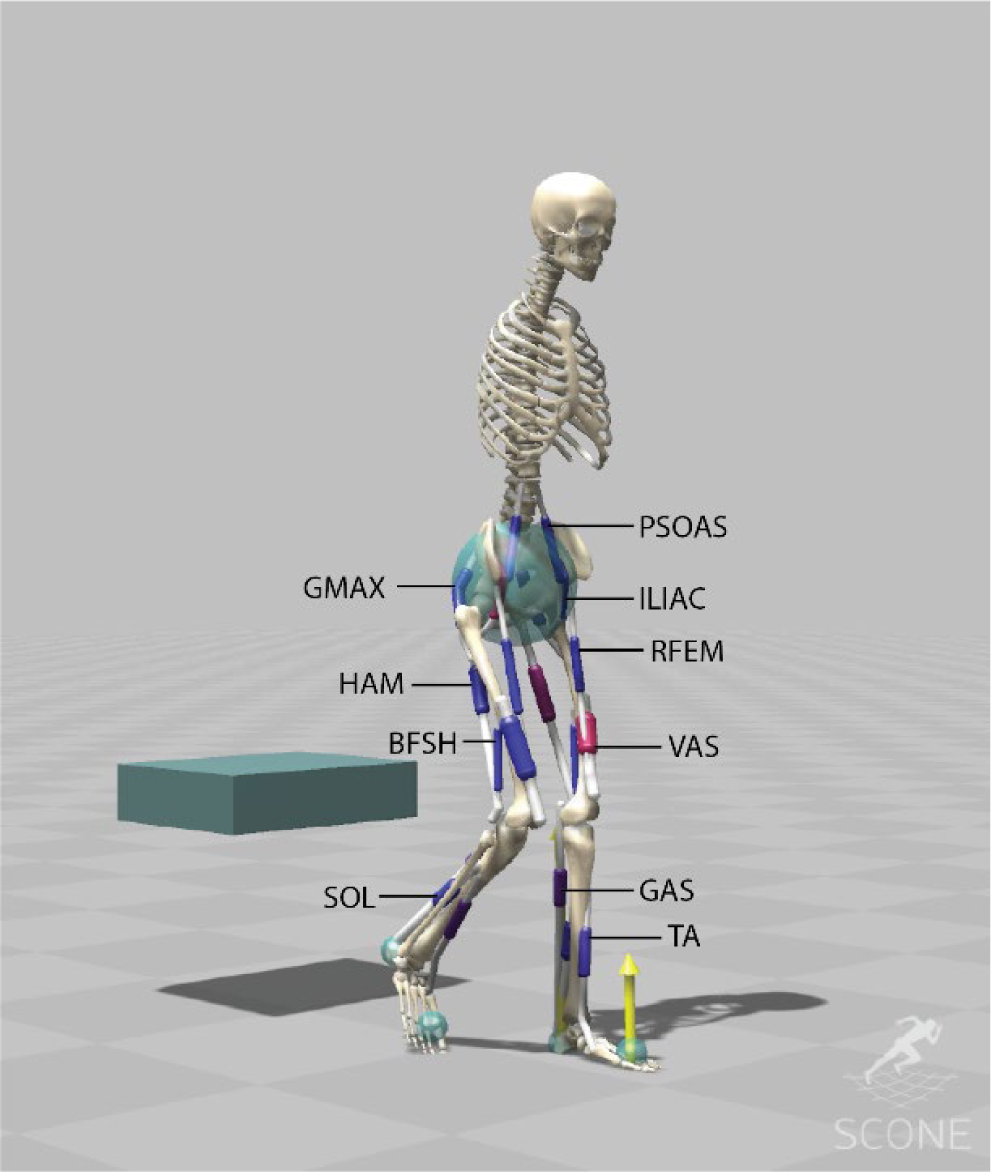
The musculoskeletal model (H1120) has 11 degrees of freedom and is actuated using 20 Hill-type muscle-tendon units. The H1120 model is available on (18), comparison with existing opensource models is provided Supporting Files Table 1 and Supporting Files Figures 1-9.

**Table 1:** Neural latencies were implemented for the monosynaptic, antagonistic and vestibular controllers as multiples of 5ms. Compared to prior publications, HAM, BFSH, and RFEM have longer latencies (15ms instead of 10ms, 20ms instead of 10ms, and 20ms instead of 10ms respectively). Vestibular delays are higher compared to prior publications that used 20ms delays(11).

| Muscle | Monosynaptic control $\Delta_t$ (ms) | Antagonists | Antagonists control $\Delta_t$ (ms) | Vestibular control $\Delta_t$ (ms) |
| --- | --- | --- | --- | --- |
| HAM | 15 | RFEM, VAS | 20 | 45 |
| BFSH | 20 | RFEM, VAS | 20 | 45 |
| GMAX | 10 | ILIAC, PSOAS | 10 | 40 |
| ILIAC | 10 | GMAX | 10 | 40 |
| PSOAS | 10 | GMAX | 10 | 40 |
| RFEM | 20 | HAM, BFSH | 20 | 45 |
| VAS | 20 | HAM, BFSH | 20 | 45 |
| GAS | 35 | TA | 35 | 55 |
| SOL | 35 | TA | 35 | 55 |
| TA | 35 | GAS, SOL | 35 | 55 |
| Lumbar |  |  |  | 35 |
| Thoracaic |  |  |  | 30 |

**Table 2:**
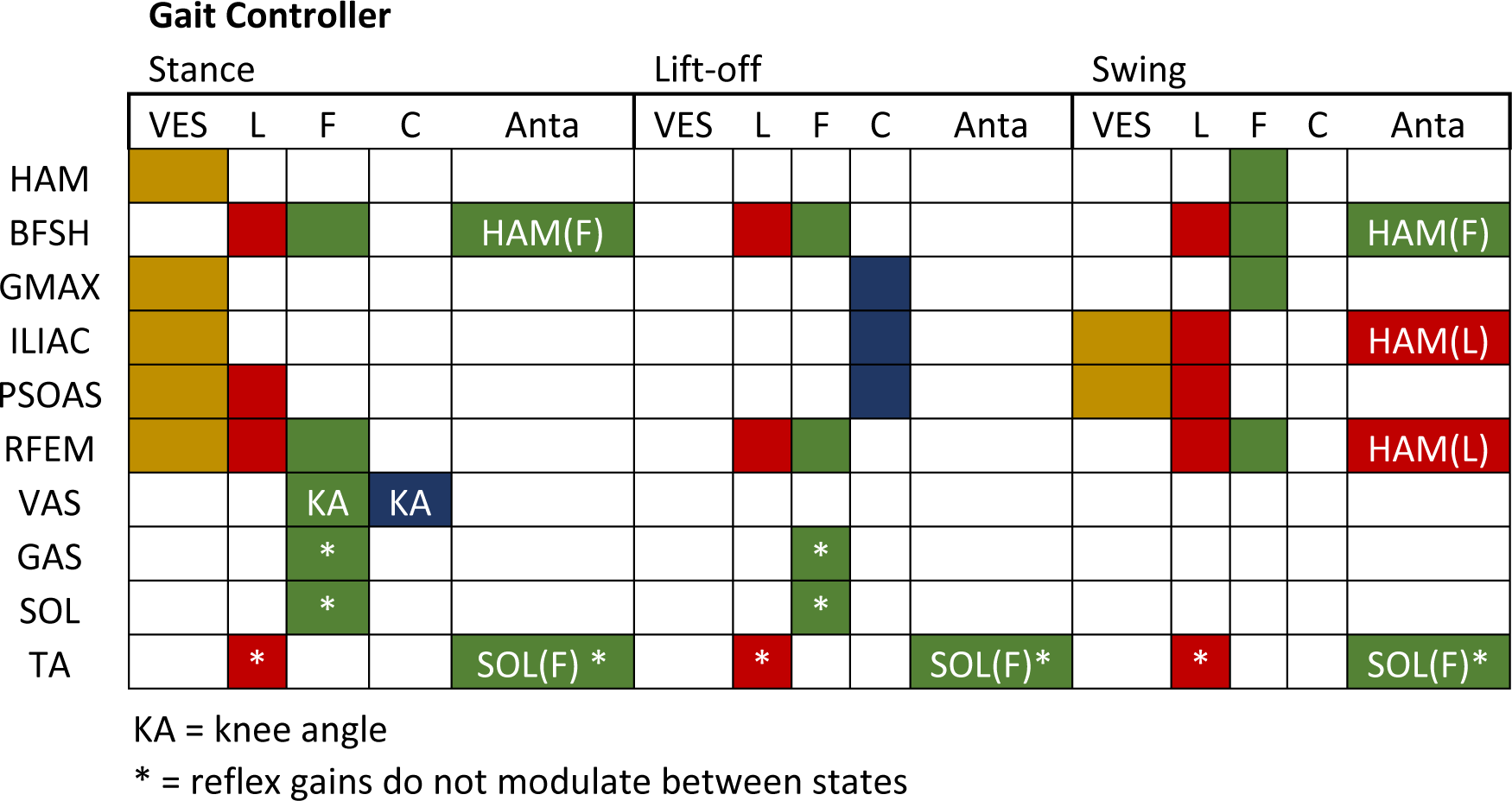
The muscle-reflex gait controller is based on a previously published controller by Geyer & Herr (2010). We added the BFSH and RFEM controller to have both uni-articular muscles around the knee and bi-articular muscles surrounding the knee and hip, respectively. Additional we added controllers for the ILIAC and PSOAS muscles. Muscle feedback pathway of the length feedback (L), Force feedback (F), and constant (C). All monosynaptic muscle-based feedback laws were positive feedback laws for length (L+). All antagonistic muscle pair-based feedback laws (Anta) were negative feedback laws for length (L-).

The model is actuated using 20 Hill-type muscle-tendon units (MTUs) (21). This reduction in number of muscles was done by combining muscles with similar functions in the sagittal plane into single MTUs with combined peak isometric forces to represent 11 major bi-lateral muscle groups: gluteus maximus (GMAX), biarticular hamstrings (HAM), iliacus (ILIAC), psoas (PSOAS), rectus femoris (RF), vasti (VAS), Biceps Femoris short head (BFSH), gastrocnemius (GAS), soleus (SOL), tibialis anterior (TA).

Peak isometric forces were based on a lower limb model of Delp et al. (1990) (Gait2392) (22), peak force from this 43-MTU model were summed to obtain a model with reduced number of muscles (Supporting Files Table 1). The optimal fiber lengths and tendon slack lengths are based on Gait2392 (23), except for HAM, BFSH, ILIACUS, PSOAS, where the updated model of Rajagopal et al. (2016) (RAJAG) was used (24). The model is free from wrapping surfaces, instead muscle paths were optimized using via-points to match the muscle moment arm data, peak normalized fiber length, and normalized path lengths (muscle-tendon lengths – tendon slack length) from existing models and experimentally measured moment arms. The lumbar and thoracic joints are torque driven. The H1120 model is available on (18), comparison with existing opensource models (Gait2392 (23), Gait1018 (25), RAJAG (24), ARNOLD (26)) is provided in Supporting Files Table 1 and Supporting Files Figure 1-9.

#### 2.1.1 H1120_STW

Contact forces between the feet and the ground and between the buttocks and the chair are modelled using Hunt Crossley force spheres and box respectively (27,28) (Fig. 2). Each foot has two contact spheres representing the heel and toes, each with a radius of 3 cm. The pelvis has one contact sphere of 12 cm representing the buttocks. The chair is represented as a box (40×12×5 cm). The contact spheres at the feet had a plane strain modulus of 17500 N/m^2^ and at the chair 10000 N/ m^2^. All contact spheres have a dissipation coefficient of 1 s/m, static friction and dynamic friction coefficients of 0.9 and 0.6 respectively. Joint forces have a limit stiffness of 500 N/m, representing the ligaments.

#### 2.1.2. Hyfydy model

The model was developed first as an OpenSim 3 model and translated into a Hyfydy model to improve the simulation speed (20). Both the OpenSim and Hyfydy version of the model are publicly available.

### 2.2 Controllers

In the development of the sit-to-walk controller, we have tested several combinations of controllers, using muscle-reflex controllers and existing gait controllers (13). The most realistic and stable controller was a combination of two reflex-controllers, which is in line with what other research groups found for the sit-to-stand movement (15), followed by gait control.

#### 2.2.1 Stand-up controller

The stand-up controller is based on a reflex controller with two phases, each containing its own set of control parameters. The reflex controller is based on monosynaptic and antagonistic proprioceptive feedback from the muscles and vestibular feedback linked to the pelvis tilt (Table 1, Fig. 3). The proprioceptive control is described by:

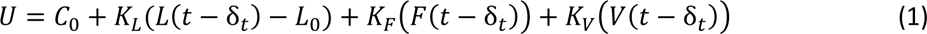

**Figure 3:**
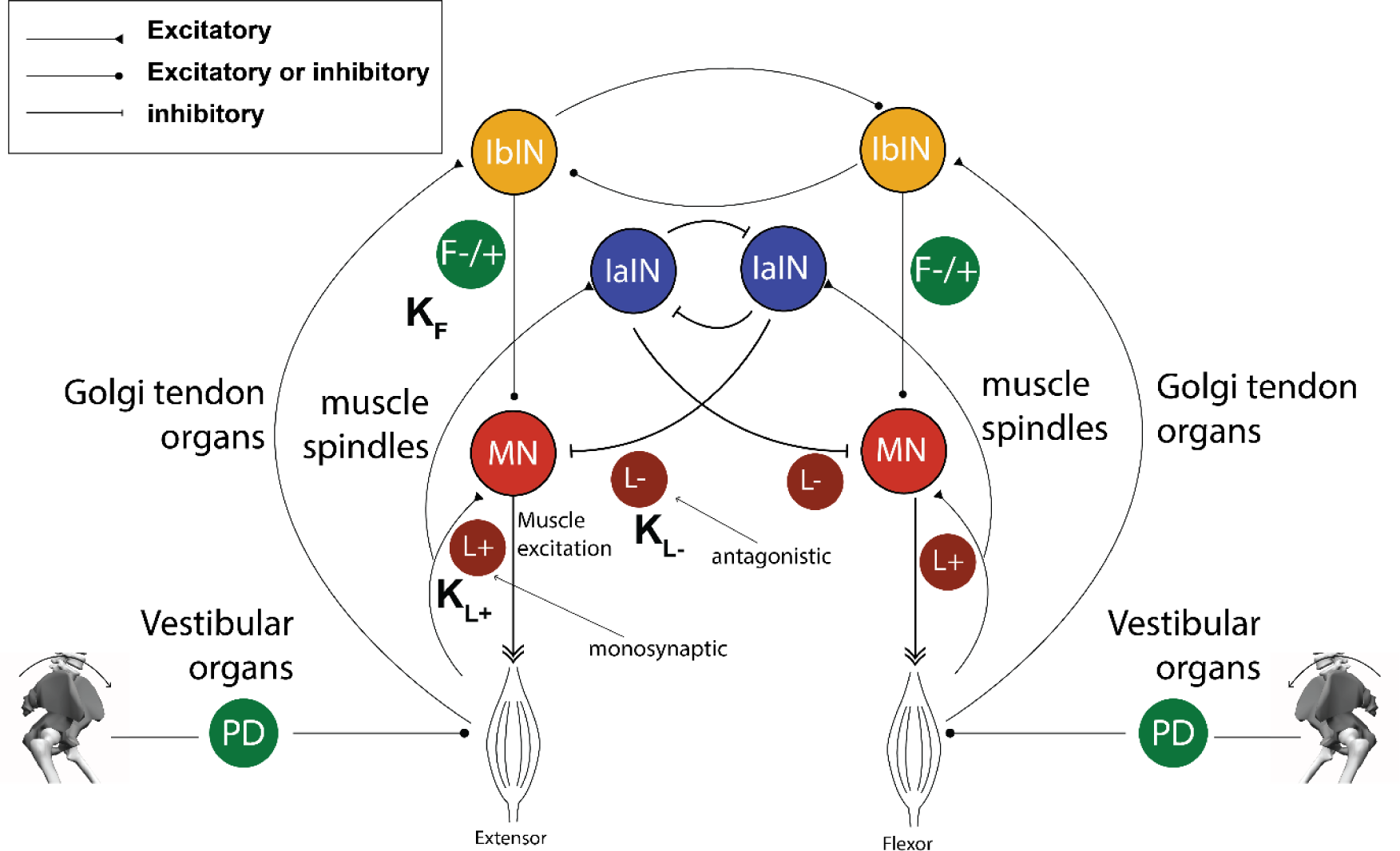
The reflex controller is based on proprioceptive and vestibular feedback for each muscle. Each muscle has a mono and/or antagonistic reflex control (Table 1). The standing up controller is based on two consecutive controllers (two phases).

In which *K_L_, K_F_, K_V_,* the gains of the controller, and *C*_0_, constant actuation, are optimized., *L*_0_, the length offset, is set to 1; although we could have optimized for this parameter, the result is a constant actuation (*K*_*L*_ *L*_0_) hence the optimization would not have been able to distinct this from *C*_0_. *L*(*t*) is the normalized CE length (*L* /*L*_*opt*_) at time *t*. The torque driven lumbar and thoracic joints were controlled by a PD control and were dependent on the velocity and position of the respective joints.

##### Delays

Neural latencies were implemented for monosynaptic, antagonistic and vestibular feedback as multiples of 5ms due the range of the accuracy we could obtain for these parameters from literature. For the monosynaptic controllers for SOL the delay was set to 35ms, based on the experimental measurements of the mean H-reflex latency (30.2ms, SD = 2.1), plus the distal motor latency (3.3ms, SD = 0.6ms) (16,29). The latency of the reflex loop for GAS and TA were also set to 35ms with the assumption of a similar length of reflex loop. RFEM latency was set to 20ms, based on the patellar tendon reflex latencies (21ms, SD = 1.5ms) (29) assuming the muscle spindle latency to be negligible (30). We assumed that VAS, and BFSH have a similar neural path length (31,32). We do expect the VAS and BFSH to have a slightly higher latency than the RFEM, but within the 5ms margin. HAM has a lower latency and was set to 15ms.

The other latencies were estimated based on the difference in motor distance between SOL and the particular muscle and the motor axon conduction velocity (CV) (about 50 m/s). We estimated the difference in motor distance between GMAX and SOL to be 50-60 cm, which resulted in 10ms latency for GMAX. We assumed PSOAS and ILIAC to have a similar path length as GMAX from the spinal cord. For the antagonistic controllers, we took the average of the delays of the two muscles involved.

The vestibular delay is based on electromyographic responses following galvanic vestibular stimulation (33). Note that large inter-and intra-subject differences have been reported on these responses (34). In general, the vestibular latency is a summation of latencies within the neural system, based on conduction velocity (CV) and the central nervous system (CNS) distance:

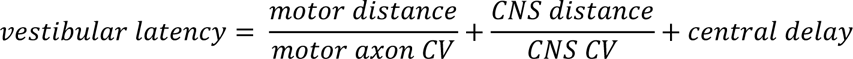

We looked at the short latency responses. SOL has a mean vestibular short latency delay from stimulation to activation onset of 55ms (33,35). The latencies of the responses in TA are equivalent to those in SOL (35) and we assumed the GAS to have a similar response time. H-reflex latency plus the distal motor latency of SOL represent the time to travel the motor distance twice, plus the neuron delay in the spinal cord. With these values of SOL we could make a rough estimate for the other muscles on the vestibular latencies (Table 1):

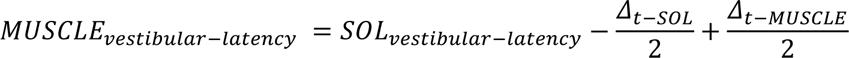

The values were round down to integers of 5 ms. Results are in line with inter-muscular differences in medium latency delays reported in literature (34). For the lumbar joint we used 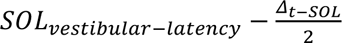 (35 ms). For the thoracic joint, we used an estimate of 30 ms based on the central delay and CNS CV estimates from (34).

#### 2.2.2 Gait controller

The gait controller is based on a previously published controller by Geyer & Herr (2010) (11). We added the BFSH and RFEM to the Geyer & Herr (2010) controller to have both monoarticular muscles around the knee and bi-articular muscles surrounding the knee and hip, respectively. Both muscles are controlled through monosynaptic proprioceptive length and force feedback paths, with separate gains for each of the three states. For RFEM we added vestibular feedback control in Stance. We also had two additional hip flexors (ILIACUS and PSOAS) and two torque driven joints (thoracic and lumbar). We configured the ILIAC and PSOAS controllers like the HFL paths from Geyer & Herr. The torque driven lumbar and thoracic joints were controlled by a PD control (vestibular control) and were dependent on the velocity and position of the respective joints and independent of states. A PD controller was used to control the pelvis tilt angle (*θ*) and velocity (θ) with respect to ground, comparable to vestibular feedback, using the muscles crossing the hip joint (HAM, GMAX, ILIAC, PSOAS, RFEM) Note that the vestibular and neural latencies were updated to the values of Table 1 and therefore differ from previous models.

#### 2.2.3 Sit-to-walk controller

The sit-to-walk controller is a sequential controller consisting of the standing up controller (two-phase reflex controllers) and a gait controller. The transition times between the reflex controllers and the gait controller were free parameters included in the optimization process.

States calculated from the model were muscle length (*l*), muscle velocity (*v*), muscle force (*F*), and pelvis tilt orientation (*θ*) and velocity (θ). In total, there were 551 free parameters of the optimization, including the controller gains (*K_C_*, *K_L+_*, *K_F±_*, *K_p_*, and *K_v_*), offsets for muscle length feedback (*l_o_*), proportional feedback of *θ* (*θ_o_*), proportional feedback of the lumbar and thoracic joints, transition times between the controllers, and stance load threshold for the gait state controller.

### 2.4 Simulation and optimization framework

The equation of motion and their integration was done in SCONE (36). The simulations started with different initial postures of the model’s dof, while the velocities were set to 0. The selected initial positions were based on commonly used seating positions prior to stand-up (Fig. 4). The simulation stopped when the maximum simulation time (12s) was reached or until the model fell, which was defined as the COM height being below 0.6 times of its height at the start of the simulation.

**Figure 4:**
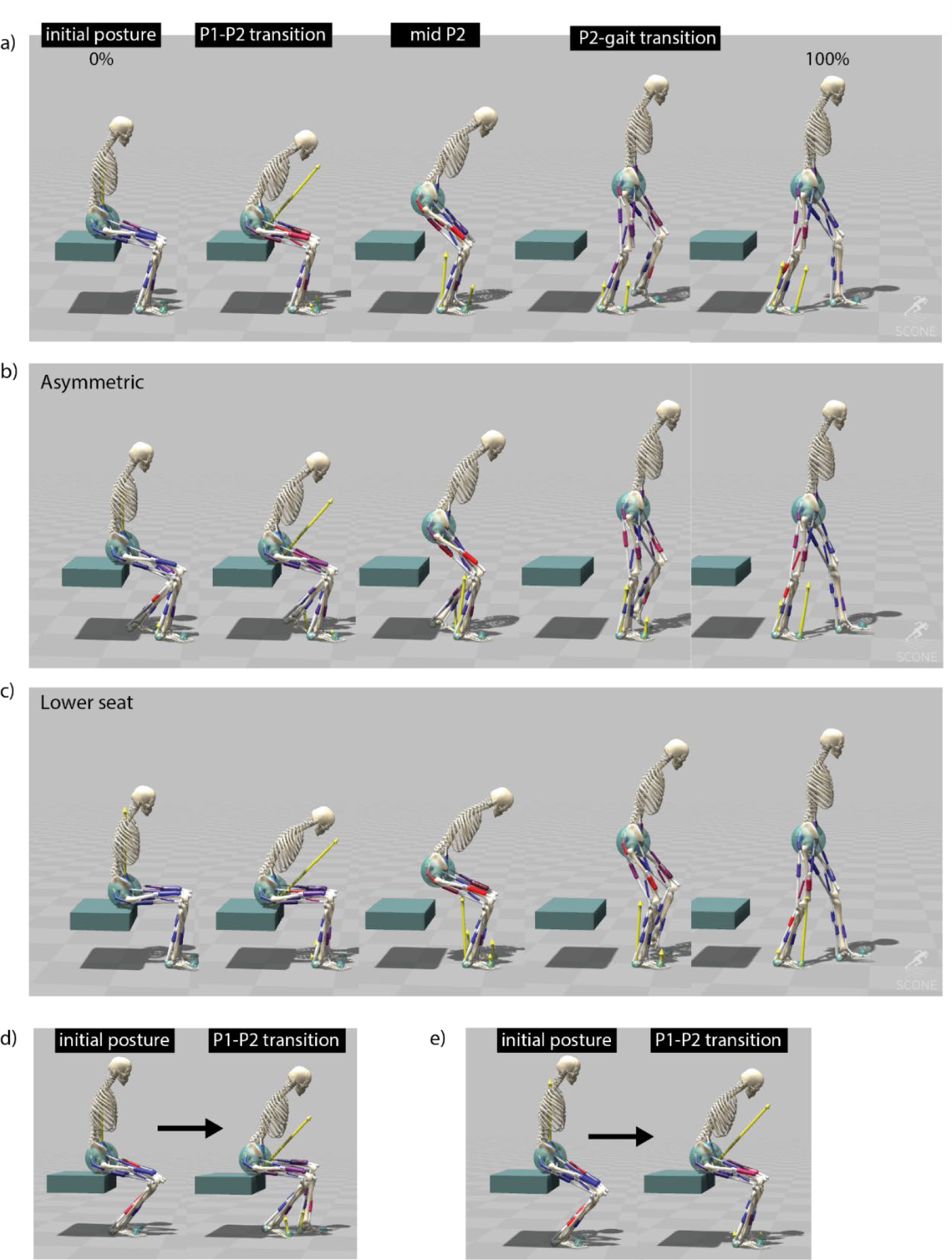
Sit-to-walk simulations; a) simulation with normal seat (seat height 44cm). Shown are the postures at t = 0, at the transition between the reflex controllers (P1-P2), midway P2, and at the transition from P2 to gait control; b) simulation where the initial posture of the model is an asymmetric position in the anterior-posterior direction; c) simulation with a lower seat height (35 cm); d,e) initial posture and posture at end of the first controller for a lower seat (d) and a higher seat (e). The initial feet positioning of the model was set backwards. The model uses P1 to reposition the feet to a better starting position for sit-to-stand transitioning.

The optimization objective was to minimize the weighted sum of a set of measure terms, each of contributing to a high-level description of the sit-to-walk task.

#### Mimic measure

To save simulation time and increase optimization performance, a measure was used that defines how well the simulation mimics measured COM pelvis position (x, y). The purpose of this measure was to have a rough standing up trajectory for the model to start from without dictating any specific motion pattern; This measure becomes zero once the motion is roughly withing the boundary of the trajectory.

#### Gait velocity measure

This measure computed the average forward velocity of the model, and was used to promote locomotion at a predefined minimum velocity of 0.8 m/s. This speed was considered the minimum required speed to have a gait pattern that represents gait inside the lab and clinic. In daily life, normal gait speeds for older adults typically range around 0.9 m/s for women and 1.0 m/s for men (37). As such, a speed of 0.8 m/s was considered a realistic minimum for measurements conducted in clinical environments. The value is normalized by the set minimum velocity. To allow the model time to reach the minimum velocity after standing up, the first three steps were not considered in the gait velocity evaluation. This measure becomes zero once the minimum velocity criterion is met.

#### Range measure

These measures penalize excessive joint limit forces and range of motion from ligaments, soft tissue, and joint geometry that are not present in the simplified model: lumbar extension (−50…0 degrees), thorax extension (−15…15 degrees), pelvis tilt (−50…30), ankle angle (−60…60). For the knee, there was a penalty on the DOF limit force (500 N/m) was out of range. All range measures become zero once within range or below their given thresholds.

#### Minimization of head acceleration

If no minimization for the head acceleration was set, simulation would produce a large initial peak in the ground reaction forces in gait. To represent how humans stabilize their vestibular and visual systems during gait, we added a head acceleration penalty with a threshold of 1m/s^2. This penalty measure returned to zero once the head acceleration is below the threshold.

*Energy objective:* Finally, an energy estimate was used as to minimize energy consumption during stand-up and minimize cost-of-transport during gait (40) The measure consisted of metabolic energy expenditure based on (38,39) (*J*_*mb*_), and was complemented by the minimization of cubed muscle activations (*J*_*act*_), along with torque minimization ( *J*_T_) at the lumbar and thoracic joints to serve as a proxy for the absence of trunk muscles. During the optimization process, the overall energy estimate, a weighted sum between the terms (*J_total_ = w_mb_J_mb_ + w_mb_J_act_ + w_T_J_T_*) became the sole non-zero term within the objective function. We empirically set W_*mb*_= 0.01, W_*act*_ = 0.1 and W_T_= 0.0003 for all conditions.

##### 2.4.1 Optimization algorithm

Optimization was performed using the Covariance Matrix Adaptation Evolutionary Strategy (CMA-ES) in the Open Source software SCONE (36). The population size of each generation was 10, the number of iterations was dependent on the simulation time. In line with previous studies (14), we have run multiple parallel optimizations with the same initial guess and used the best set as start the next set of optimizations.

### 2.5 Altered conditions, neuromuscular capacity, and movement objectives

One of the applications for the controller is to determine how different seats or initial postures affect the sit-to-walk biomechanics and adaptation strategies. As an example, we used the controller to learn STW from a lower seat (height = 35cm) (Fig. 4a), and when the feet were placed asymmetrically (anterior-posterior position) (Fig. 4b). We refer to the leg that is in swing during the initial step as the “stepping leg,” and the leg that is in stance during the initial step as the “stance leg.” We tested several other initial postures which are available (18).

### 2.6 Experimental data

To validate the simulations, we adopted a subset of kinematic and kinetic data from van der Kruk et al. (2022) (6), where kinematic and kinetic data of sit-to-walk at self-selected speed was recorded of young 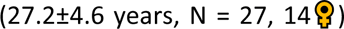 and older 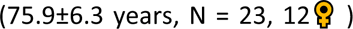 adults. This study received ethical approval by the institutional ethics committee of Imperial College London, UK. All participants gave written informed consent. Participants were seated on an instrumented chair with armrests, which was adjusted in height to have an approximate 90^0^ knee angle while seated in neutral position. The sit-to-walk movement was performed at self-selected pace and as-fast-as-possible each with 5 repetitions. The initial posture of the participants was not restricted (e.g. foot position, trunk angle, position on the seat), and participants were allowed to use their arms. For the verification purposes, only the trials of young participants that naturally stood up without arms were selected (45 trials). Note that the arms were not crossed on the chest, so arm swing was still allowed. Reaction forces were measured via two Kistler forceplates embedded in the walkway and one at the seat, and two 9129 AA Kistler forceplates in the armrests. A Vicon system with 10 cameras (MX T20) captured the STW volume (100 Hz). Participants were equipped with 84 reflective markers. A wireless Delsys EMG system with 16 sensors measured the EMG signal of specific muscle groups. The European guidelines for the EMG sensor placement were followed.

Marker trajectories and GRFs were translated into joint kinematics (inverse kinematics) and muscle forces (static optimisation) in OpenSim 4.2 using the musculoskeletal model of Rajagopal et al. (2016) (24). To compare muscle activations, a weighted sum of the muscle activations from the grouped muscles was calculated. Activations were weighted by the maximum isometric force of individual muscles divided by the total isometric force of the grouped muscles.

Muscle activation levels were compared to measured sEMG data for seven bi-lateral muscles (HAM, GMAX, RFEM, VAS, GAS, TA). Muscle excitations were determined from sEMG data that were high-pass filtered (30Hz), then full-wave-rectified, and low-pass filtered (6Hz) with zero-lag fourth order Butterworth filters. Since no Maximum Voluntary Contraction was available in the dataset, the excitation was normalized by the maximum excitation measured within the individual complete sit-to-walk trials (stand up-walk 3 meter –turn – walk 3m – sit down). Note that as a result the activation levels indicated by the sEMG might be higher than the actual activation levels.

Simulations were verified by comparing joint angles, ground reaction forces, and muscle activations.

## 3. Results

### 3.1 Sit-to-walk controller (normal posture)

#### 3.1.1 Kinematics

The sit-to-walk simulation consists of three phases (P1, P2, GAIT). During P1, the model repositions the trunk, bending it forward (Fig. 4a). This repositioning has been extensively reported in experimental research(17,40), and is in line with the measured kinematics (Fig. 5). We noted that when we changed the initial posture of the feet to a far backwards position, the P1 controller repositioned the feet, placing them symmetrically or asymmetrically forward (Fig. 4 d, e). This is beneficial for the transition of the movement into the standing up and gait phase, and shows that the model is able to find realistic foot and trunk adaptation strategies.

**Figure 5:**
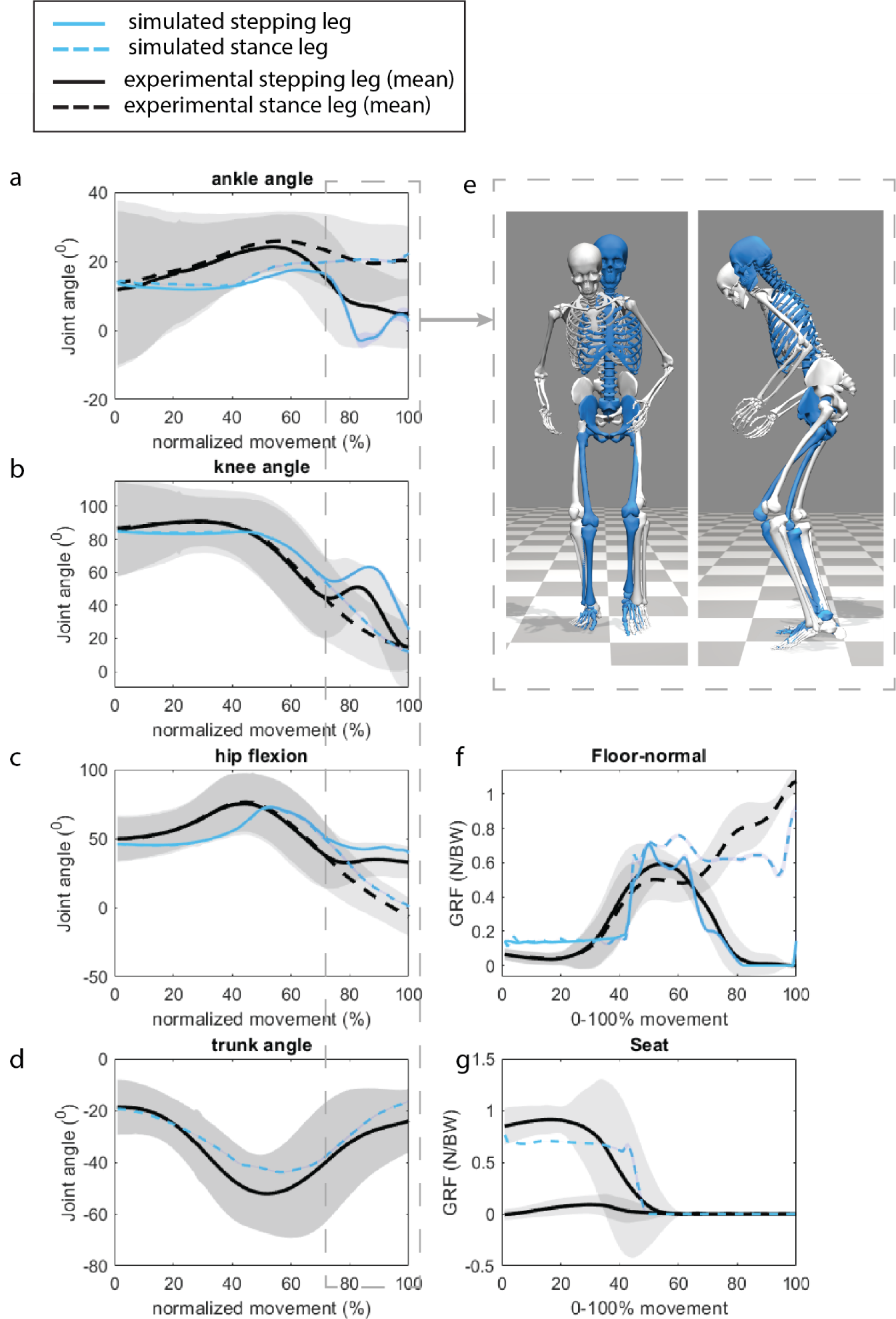
Measured and simulated joint kinematics (a-d), ground reaction forces (f), and seat reaction force s(g). The light grey areas indicate the mean±2SD of the experimental data. The increased knee and hip flexion observed during the initial step (at 80% of the motion) are attributed to the absence of pelvis rotation in the planar model (e).

The ankle dorsiflexion, knee flexion, and hip flexion exhibit a shallower trajectory in P1 when compared to the measured kinematics. This primarily arises from the simplified contact geometries of the chair and buttocks. In reality, the thighs roll over the chair’s surface, resulting in a more extended contact period, which is not accounted for in the simplified contact geometry.

In P2, the model is extending from a seated position into the GAIT controller, transitioning into gait before the joints are fully extended (Fig. 4a). This finding aligns with established principles of human movement control during the sit-to-walk transition (5), as indicated by the similarity observed in the knee and hip angles as measured (Fig 5). The increased knee and hip flexion observed during the initial step (at 80% of the motion) are attributed to the absence of pelvis rotation in the planar model, which is necessary to achieve ground clearance (Fig. 5e).

#### 3.1.2 Muscle activation

Since Maximum Voluntary Contraction was not available in the dataset, the excitation was normalized by the maximum excitation measured within the individual complete sit-to-walk trials; as a result, sEMG activation data might be overestimated, and the focus was on the time of activation and the shape of the activation curve rather than the absolute difference in activation level. We found the simulated muscle activation timing to resemble the sEMG data (Fig. 6). VAS shows a close to maximum activation, which has to do with the relatively weak VAS muscle in the H1120 model.

**Figure 6:**
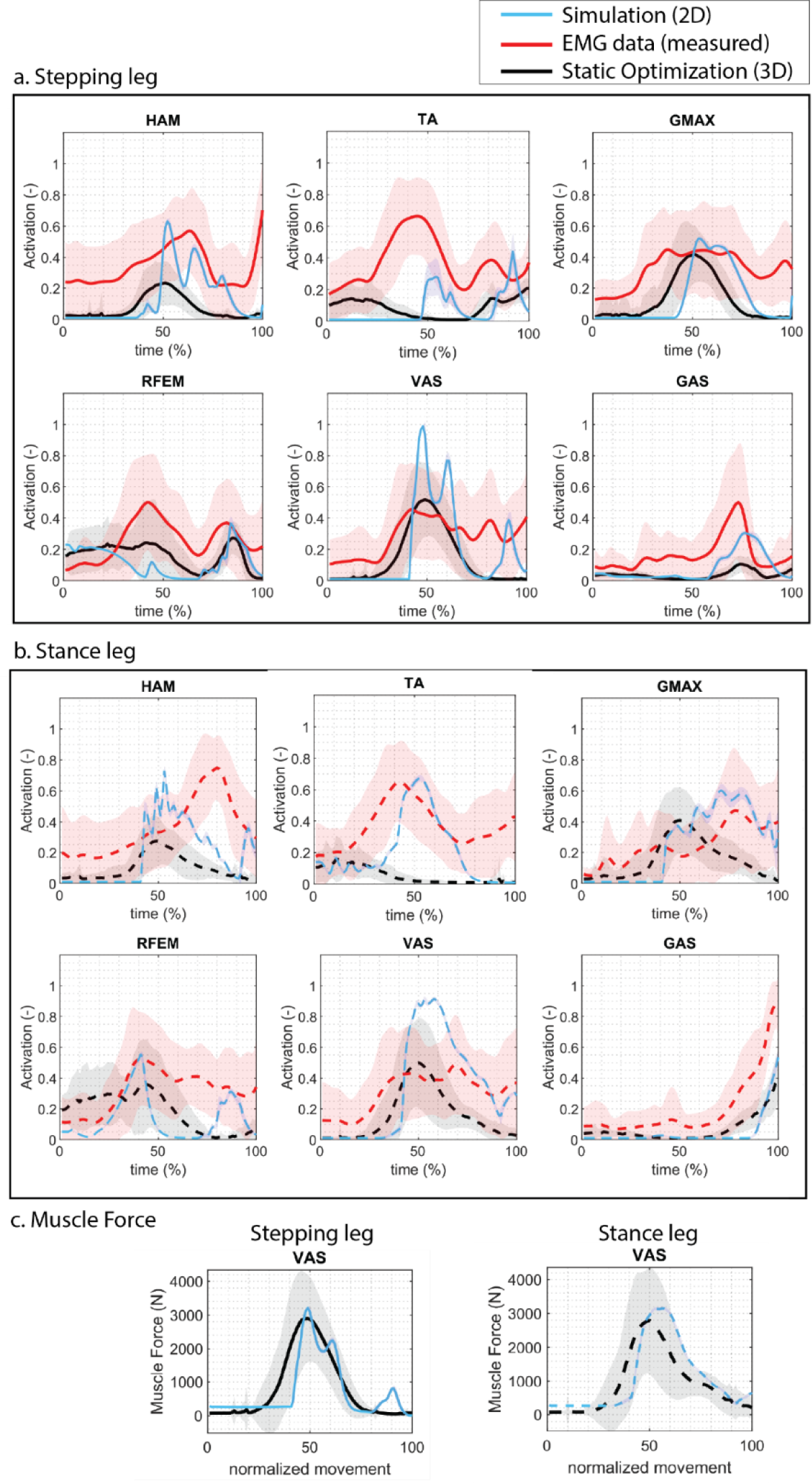
Muscle Activations. In red, the mean measured EMG data of experimental trials. Measured EMG data is normalized to the peak signal measured within the complete STW trial for each trial per participant, since no data on maximum voluntary contraction was available. Therefore, the normalized EMG data can be overestimated. In black, the mean muscle activations obtained from the inverse dynamics approach with static optimization of a 3D musculoskeletal model and measured kinetics; shade indicates +/-one standard deviation. In green, the muscle activations from the predictive simulation with the 2D controller and model.

#### 3.1.3 External forces

As a result of the simplified contact geometries of the chair and buttocks, there are differences in the ground and seat reaction forces. Simulated seat reaction forces are less smooth than ground truth seat reaction forces. The foot ground contact forces are underestimated for the stance leg in the initial step (Fig 5e).

#### 3.1.4 Joint Loading

When comparing the simulated muscle forces with those obtained through static optimization (RAJAG model (24)), the forces are comparable (Fig. 6c). The peak simulated joint loading closely aligns with static optimization for all joints, except for the ankle joint of the stance leg during the first step, where the predictive simulation underestimates the ankle joint loading (Fig. 7). This is due to the underestimation of the ground contact force of the stance foot (Fig 5f).

**Figure 7:**
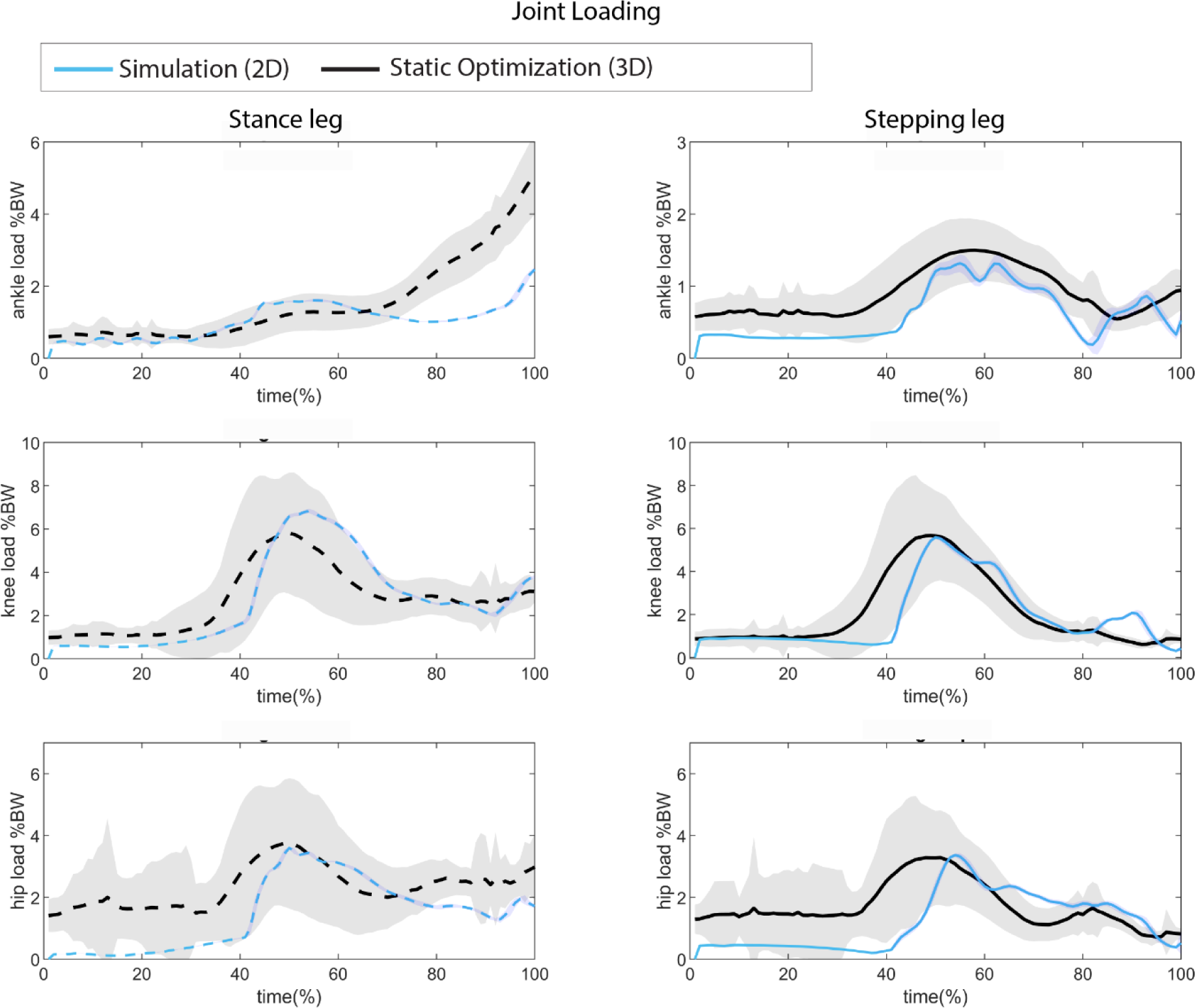
Knee joint load comparison between measured knee load estimation (inverse kinematics, static optimization, 3D musculoskeletal model (RAJAG)) and the simulated knee joint loading. The load is comparable.

### 3.2 Alternative Conditions

#### 3.2.1 Lower seat height

When the seat height was lowered, the model exhibited a greater trunk angle in P1 compared to the normal seat condition (with a peak trunk angle of 51^0^ in the low seat versus 42^0^ in the normal seat condition), aligning with principles of human motor control (5) (Fig. 4c). As expected, the lower seat necessitates increased muscle activations and leads to higher joint loads (Fig 8). The lower seat results in higher peak joint loadings compared to the normal seat: knee load +15 and +21%, ankle load +21% and +51%, hip +9% and +10% for the step and stance leg respectively. The biggest differences in muscle activation is an increased activation of the BFSH, TA, and PSOAS in the rising phase of the stepping leg, and the GMAX in the rising phase of the stance leg (Fig 8a).

**Figure 8:**
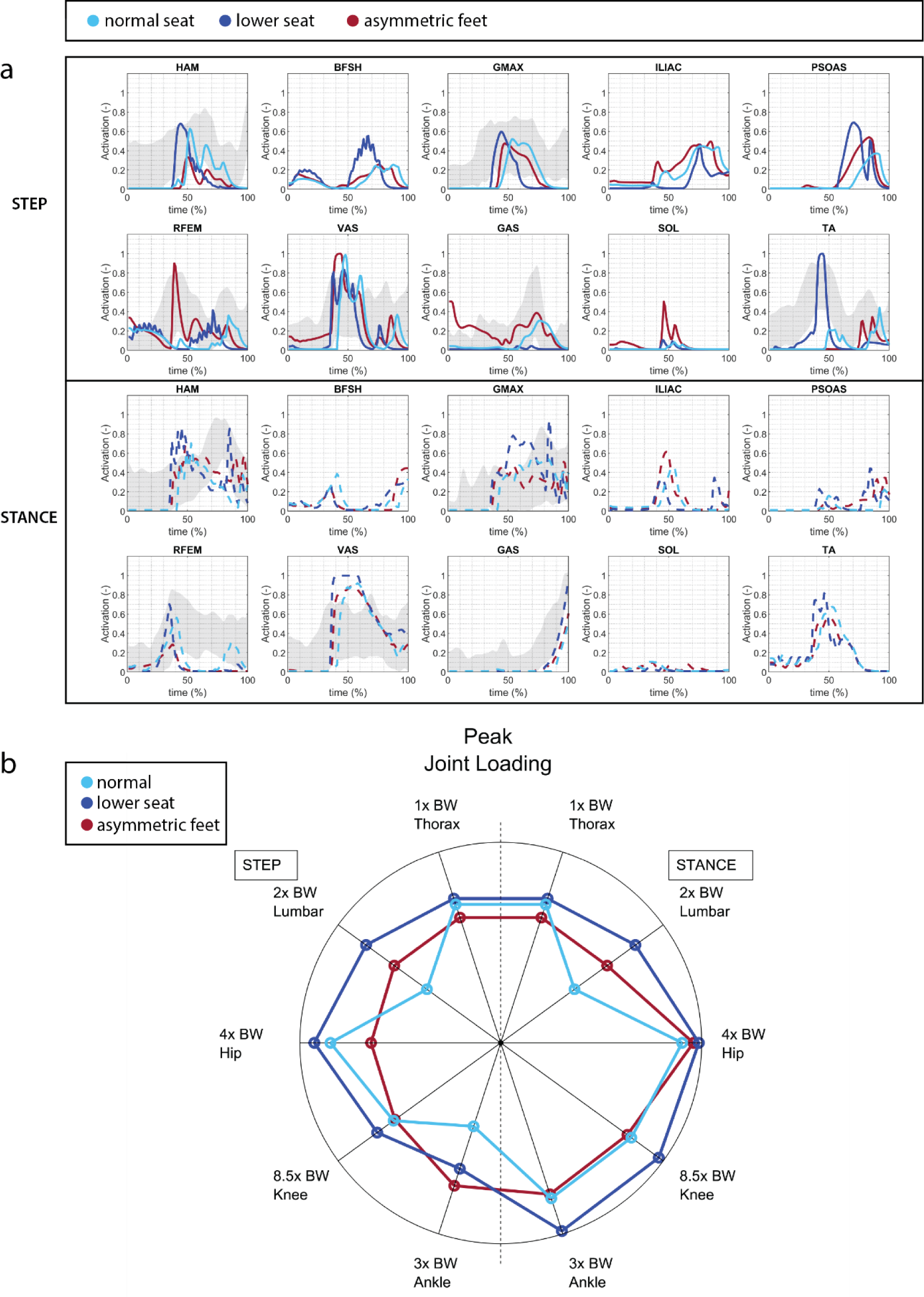
a) Muscle activation for lowered seat and asymmetric foot positioning. the lower seat requires higher muscle activations than the normal seat condition. B) peak joint loading for alternative seat height and foot positioning. Note that the thoracic and lumbar joints are torque actuated joints, so the load resulting from muscle force contraction is missing in these joints. The lower seat condition results in higher joint loads. The asymmetric foot positioning in which the stepping leg is placed backwards, leads to a large reduction of peak hip load in the stepping leg with the drawback of increased peak ankle load in the stepping leg.

#### 3.2.2 Asymmetric feet position

During asymmetric foot positioning, the trunk remained more upright compared to the symmetric condition (Fig. 4b). This posture contributes to a significant reduction in hip load in the stepping leg compared to the symmetric condition with a decrease of 24% (Fig. 8). The bilateral knee loads are also slightly reduced (−1-3%). However, at the onset of movement, the ankle load in the stepping leg is notably higher by 71% compared to the symmetric foot condition. There is less activation of the HAM in the stepping leg, while RFEM and GAS show heightened activated, with both higher peak and duration, compared to the symmetric feet position in the stepping leg. In summary, adopting an asymmetric foot positioning where the stepping leg is positioned backward results in a decrease in the peak hip load in the stepping leg. However, this comes with the trade-off of an increased peak ankle load in the stepping leg.

## 4. Discussion

Progression of neuromuscular decline can lead to incorrect or insufficient compensation strategies (maladaptation), both through altered muscle recruitment and changes in movement trajectories. Control policy based predictive neuromuscular simulations are an effective tool for discovering those strategies, enabling clinicians to design treatments that are both more effective and efficient for prolonging the mobility of the patient. In this study we have developed a planar predictive neuromuscular simulation of the sit-to-walk movement based on reflex controllers. Our model produced sit-to-walk movements that match real-world kinematic and kinetic recordings. The model accurately predicts muscle activation patterns, kinematics, and joint loading. Caution has to be taken with the ankle peak loads of the stance leg during the initial step which were underestimated. We demonstrated that the model can produce realistic compensation strategies, by changing conditions such as seat height and initial foot positioning.

We found that a 2-phase reflex based neural controller can realistically simulate the task of standing up (41), which is in line with the planar sit-to-stand simulation from Muñoz et al. (2022) (15), and is different from the commonly reported 4-phase kinematic phases (40). In our model, the transition time is part of the optimization. The model is sensitive to this timing, especially when the conditions like seat height are changed, which requires a quick re-optimization of the transition times for new conditions. Prescribing the transition as a kinematic parameter would ignore the fact that the adaptation of reflex gains occurs before a change in kinematics (42). We therefore believe it is important keep this parameter independent until a valid sensory input is determined from experimental studies (42).

Our planar sit-to-walk framework allows for asymmetric strategies, like asymmetric foot positioning, and asymmetries in muscular capacities. Previous studies on sit-to-stand simulation did not allow for asymmetry (10,15) which limits the available compensation strategies, and may not properly reflect daily life movements as a result (17). Asymmetric strategies in standing up may carry potential implications for long-term joint degeneration as they alter joint loading. Simulations can offer insights for clinicians in advising patients on optimal strategies to prevent overloading or unloading following surgery or injury. The simplified planar joints in the model do not account for the complexities of medial-lateral and internal-external rotational stability of joints. Nonetheless, the planar simulations demonstrate that asymmetric foot positioning results in a significant decrease in unilateral peak hip load on the side that is placed backwards and transitions into gait first (stepping leg).

Based on the most recent meta-analysis available, there is still no consensus on a definitive torque actuator limit for upper body joints(43). To mitigate the risk of the actuator exceeding its capacity, we chose to establish the limits at 1000 Nm. The highest joint torques observed in our simulations, immediately following seat-off in the low seat condition, were 104 Nm in the thoracic joint and 167 Nm in the lumbar joint. These values align with the range documented in experimental studies (44). Consequently, we are confident that the set actuator limits have not influenced the conclusions drawn in this article.

### 4.3 Limitations

The simplified model and controller cause several simulation artefacts, of which researchers and clinicians should be mindful when interpreting results.

- During the sit-to-walk transition, humans employ medial-lateral trunk and pelvis movements to maintain balance and ensure foot-ground clearance. These dynamics cannot be adequately captured by a planar model.
- The maximum isometric forces in the model are based on (23) in which VAS muscles are relatively weak for a healthy adult, as shown by the maximum activation of this muscle while rising from the chair. When using the simulations for comparison of changes in muscular capacity, the musculoskeletal model would benefit from better estimates of the maximum isometric forces for young and older adults, and between sexes. In musculoskeletal simulations especially the relative differences between maximum isometric forces of muscles is important, due to the nature of the optimal control cost function. We expect these to differ between age groups and sexes but a clear overview and conclusion about this on a group level is currently lacking in literature.
- The buttocks are represented in the model by a single sphere. In reality, the contact with the chair extends over a longer duration as the thighs roll off over the chair. This simplified geometric representation of contact spheres leads to a flatter trajectory of ankle dorsiflexion, knee flexion, and hip flexion prior to seat-off. The stiffness and damping between the buttocks and the chair is approximately half that between the feet and the floor. We have explored and tested various ranges of values. The stiffness should be lower than that of the feet, while there shouldn’t be a ‘bouncing’ effect, which occurs when the stiffness and damping are set too low. Despite our efforts, we were unable to locate reliable experimental data for validating this specific value. In general, we expect there to exist a large variety in stiffness and damping based on both the subject and the type of chair used. While this is potentially important, we consider an exhaustive study to be beyond the scope of this paper. With respect to the foot contact model, previous studies have shown the importance of choice in foot stiffness for 3D predictive simulations (45). For our planar model we found that the simulation produced consistent results across a wide range of stiffness settings.
- Optimizations within the framework can yield various movement strategies originating from identical scenarios. In this manuscript, we opted to adopt the strategy with the lowest cost. Nonetheless, there were movement strategies closely approaching the minimal cost with different, yet equally plausible, movement strategies. In future studies, we may consider treating these simulations akin to intra-trial differences among participants and report the distribution in kinematics and kinetics, mirroring how we would handle experimental data. This approach would also mitigate the sensitivity of results to the cost function.
- Lastly, our experimental study revealed that over 80% of older adults utilized their arms to push off while standing up in a daily life setting (6). These compensatory strategies were not accounted for in this model. Introducing torque or muscle-driven arms into the model could facilitate the examination of whether this strategy is employed to compensate for diminished muscular capacity, enhance stability, or alleviate pain.

### 4.4 Conclusion

We presented a novel predictive control strategy for simulating the sit-to-walk movement. The sit-to-walk movement is a crucial aspect of the timed-up-and-go (TUG) test used in clinical assessments. Our framework is capable of simulating the effects of altered foot and seat conditions. The parts of the neuromuscular system and altered movement priorities that are prone to age-related decline are part of the framework (elements that can be adopted are depicted in Figure 1), which opens the opportunity to study the effect of age-related decline and external perturbations on compensation strategies in standing up (12,46,47).

## Data Availability statement

The source code and data for the predictive framework and the results presented in this manuscript are available from Git repository: (18).

