## Supporting Files for "A planar neuromuscular controller to simulate compensation strategies in the sit-to-walk movement"

*Table 1 – Comparison of model parameters of the H1120 musculoskeletal model with commonly used OpenSim models: G2392 (*(1)*) , G1018 (*(2)*), Rajag* (3)*, Arnold (*(4)*)*

| | | Max. Isometric Force (N) | | | | | Optimal Fiber Length (m) | | | | | Tendon Slack Length (m) | | | | | Pennation angle (rad) | | | | |
|---|---|---|---|---|---|---|---|---|---|---|---|---|---|---|---|---|---|---|---|---|---|
| | | *G2392* | *G1018* | *RAJAG* | *ARNOLD* | *H1120* | *G2392* | *G1018* | *RAJAG* | *ARNOLD* | *H1120* | *G2392* | *G1018* | *RAJAG* | *ARNOLD* | *H1120* | *G2392* | *G1018* | *RAJAG* | *ARNOLD* | *H1120* |
| Biceps Femoris Short head | BFSH | **804** | 804 | 557 | 705 | **804** | 0.173 | 0.173 | **0.11** | 0.098 | **0.11** | 0.089 | 0.089 | 0.106 | 0.322 | 0.1 | 0.401 | 0.401 | 0.264 | **0.215** | **0.215** |
| Gastrocnemius | GAS | | 2500 | | | 2241 | | 0.06 | | | **0.051** | | 0.39 | | | 0.392 | | 0.297 | | | **0.173** |
| *Gastrocnemius lateralis* | | 683 | | 1575 | 437 | | 0.064 | | 0.059 | 0.053 | | 0.38 | | 0.376 | 0.356 | | 0.14 | | 0.21 | 0.209 | |
| *Gastrocnemius medialis* | | 1558 | | 3116 | 606 | | 0.06 | | **0.051** | 0.059 | | 0.39 | | 0.399 | 0.382 | | 0.297 | | 0.166 | **0.173** | |
| Gluteus Maximus | GMAX | | 1944 | | | 1944 | | 0.147 | | | **0.147** | | 0.127 | | | 0.127 | | 0 | | | **0** |
| Gluteus Maximus (superior) | | 573 | | 984 | 109 | | 0.142 | | 0.147 | 0.024 | | 0.125 | | 0.049 | 0.039 | | 0.087 | | 0.354 | 0.382 | |
| Gluteus Maximus (middle) | | 819 | | 1406 | 546 | | 0.147 | | **0.157** | 0.147 | | 0.127 | | 0.068 | 0.05 | | 0 | | 0.367 | **0.382** | |
| Gluteus Maximus (inferior) | | 552 | | 948 | 781 | | 0.144 | | 0.167 | 0.157 | | 0.145 | | 0.07 | 0.073 | | 0.087 | | 0.382 | 0.382 | |
| Hamstrings | HAM | | 2700 | | | 2594 | | 0.109 | | | **0.098** | | 0.326 | | | 0.325 | | 0 | | | **0.202** |
| *Biceps Femoris Long head* | | 896 | | 1313 | 324 | | 0.109 | | **0.098** | 0.106 | | 0.326 | | 0.325 | 0.043 | | 0 | | 0.176 | **0.202** | |
| *Semimenbranosus* | | 1288 | | 2201 | 114 | | 0.08 | | 0.069 | 0.403 | | 0.359 | | 0.348 | 0.11 | | 0.262 | | 0.255 | 0.264 | |
| *Semitendinosus* | | 410 | | 591 | 1163 | | 0.201 | | 0.193 | 0.069 | | 0.256 | | 0.247 | 0.348 | | 0.087 | | 0.241 | 0.225 | |
| Iliopsoas | | | 2342 | | | | | 0.1 | | | | | 0.16 | | | | | 0.14 | | | |
| *Iliacus* | ILIAC | 1073 | | 1021 | 137 | 1073 | 0.1 | | **0.107** | 0.228 | **0.107** | 0.1 | | 0.096 | 0.166 | 0.093 | 0.122 | | **0.28** | 0.25 | **0.28** |
| *Psoas* | PSOAS | 1113 | | 1427 | 296 | 1113 | 0.1 | | **0.117** | 0.026 | **0.117** | 0.16 | | 0.1 | 0.115 | 0.197 | 0.14 | | **0.216** | 0.187 | **0.216** |
| Rectus Femoris | RFEM | 1169 | 1169 | 2192 | 254 | 1169 | 0.114 | 0.114 | **0.076** | 0.054 | **0.114** | 0.31 | 0.31 | 0.448 | 0.024 | 0.305 | 0.087 | 0.087 | 0.217 | **0.243** | **0.087** |
| Soleus | SOL | 3549 | 5137 | 6195 | 302 | 3549 | 0.05 | 0.05 | **0.044** | 0.193 | **0.044** | 0.25 | 0.25 | 0.277 | 0.245 | 0.243 | 0.436 | 0.436 | 0.381 | **0.494** | **0.494** |
| Tibialis Anterior | TA | 905 | 3000 | 1227 | 155 | 1759 | 0.098 | 0.098 | **0.068** | 0.095 | **0.098** | 0.223 | 0.223 | 0.24 | 0.45 | 0.223 | 0.087 | 0.087 | 0.195 | **0.168** | **0.087** |
| Vasti | VAS | | 5000 | | | 4530 | | 0.087 | | | **0.099** | | 0.136 | | | 0.115 | | 0.052 | | | **0.253** |
| *Vastus Intermedius* | | 1365 | | 1697 | 906 | | 0.087 | | **0.099** | 0.038 | | 0.136 | | 0.202 | 0.282 | | 0.052 | | 0.063 | **0.079** | |
| *Vastus Lateralis* | | 1871 | | 5149 | 1024 | | 0.084 | | 0.099 | 0.099 | | 0.157 | | 0.221 | 0.106 | | 0.087 | | 0.253 | 0.321 | |
| *Vastus Medialis* | | 1294 | | 2748 | 2255 | | 0.089 | | 0.097 | 0.099 | | 0.126 | | 0.2 | 0.13 | | 0.087 | | 0.422 | 0.517 | |

### MUSCLE MOMENT ARMS

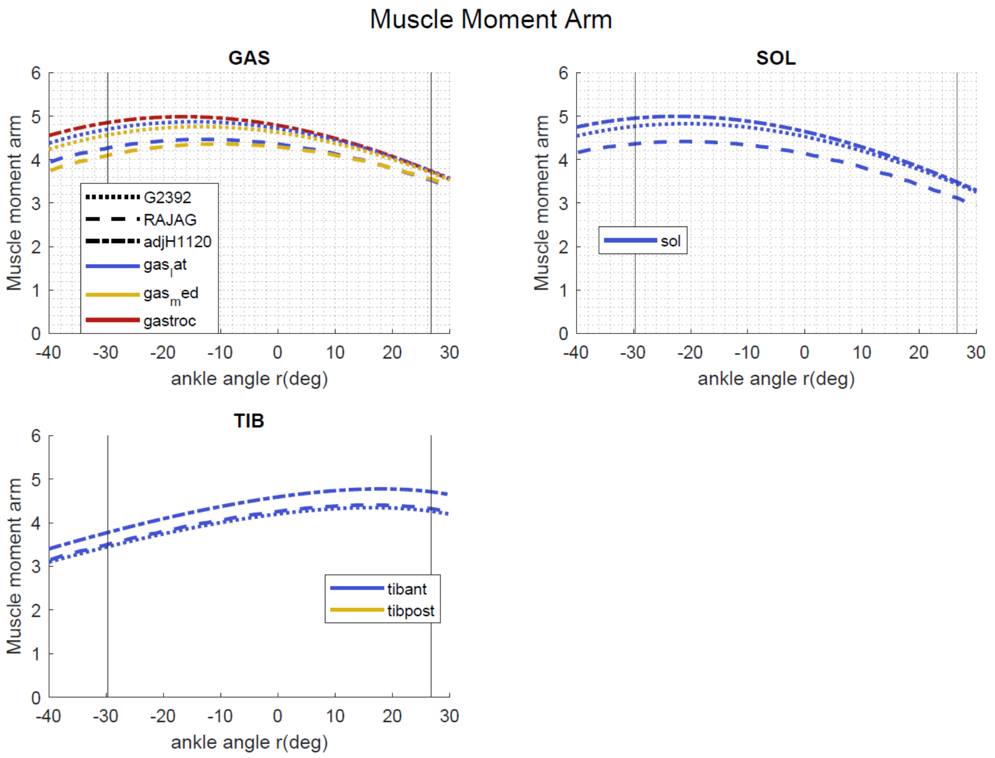


Figure 1 – Muscle Moment arms ankle joint. All other joints are in neutral (extended) positions

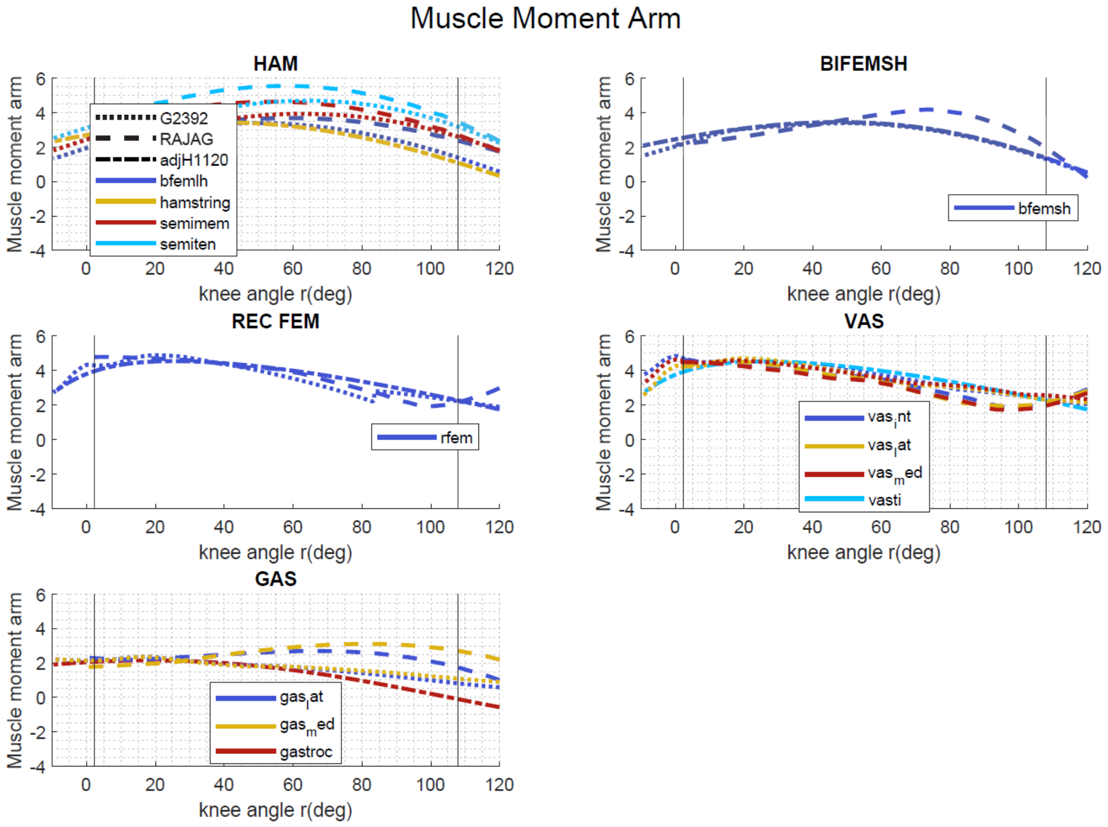


Figure 2 – Muscle Moment arms knee joint. Other joints are in neutral positions.

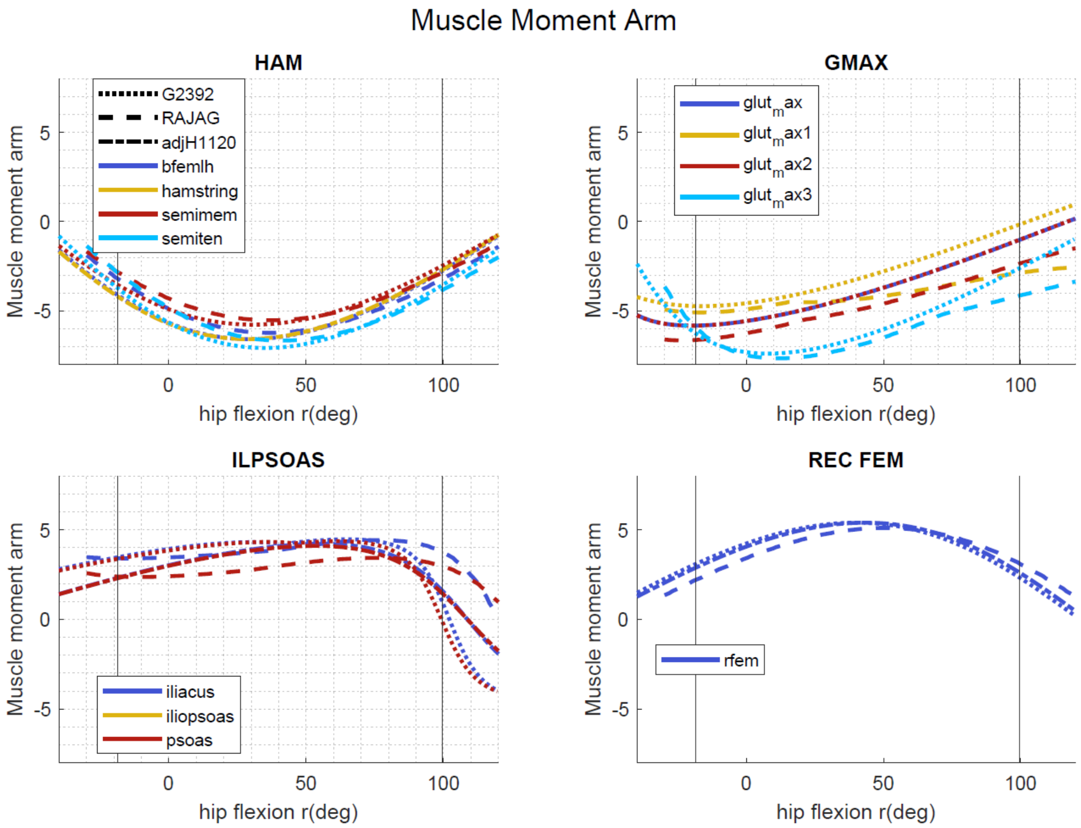


Figure 3 – Muscle Moment arms hip joint. Other joints are in neutral positions.

#### NORMALIZED FIBER LENGTHS

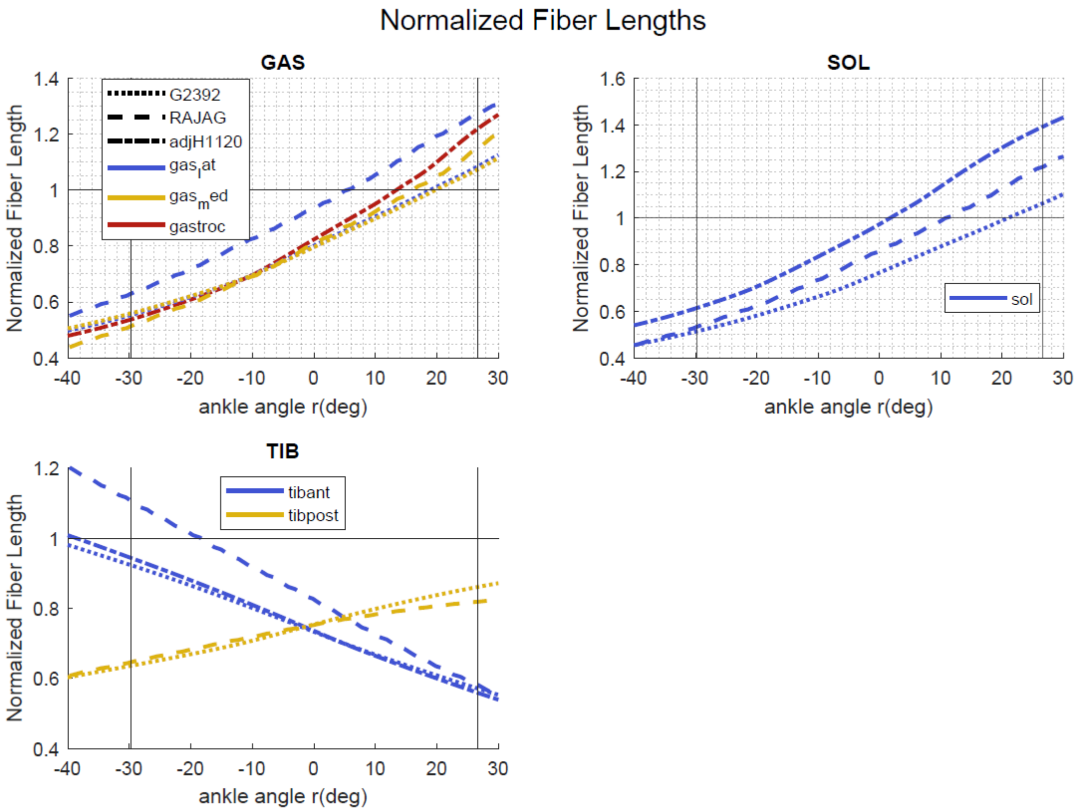


Figure 4 – Normalized Fiber Lengths of the ankle joint. Other joints are in neutral position.

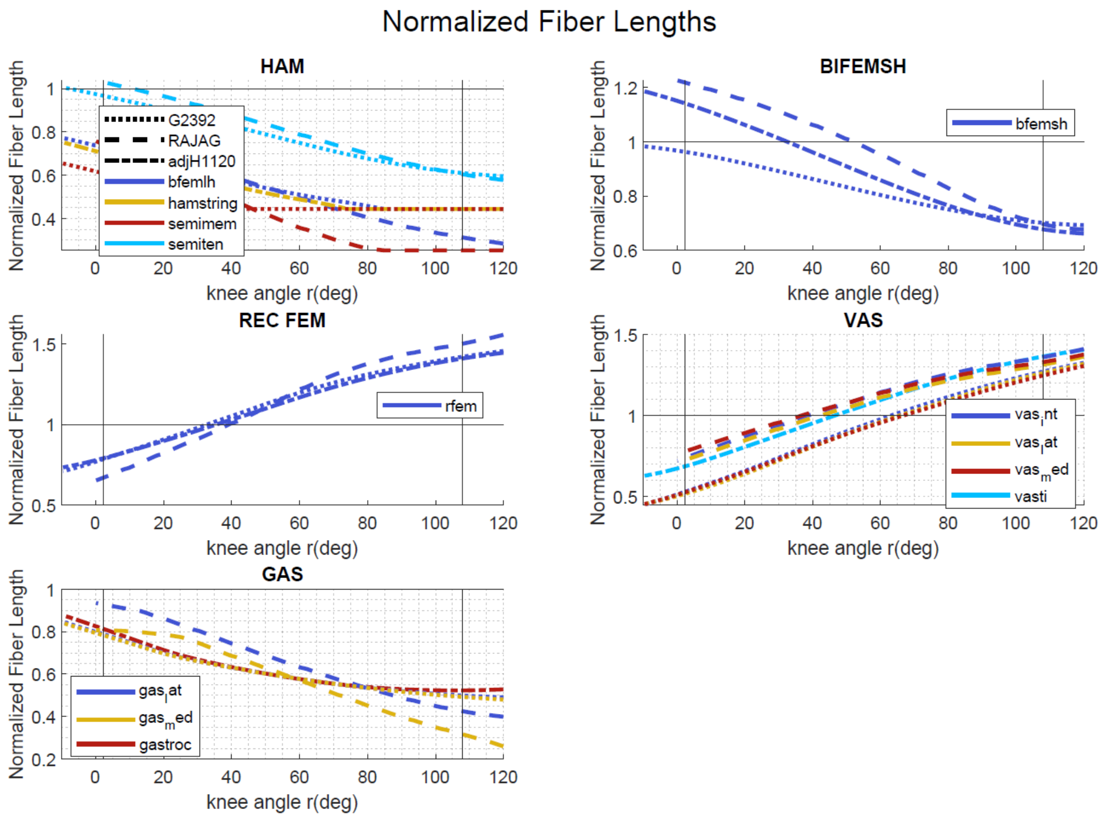


Figure 5 – Normalized Fiber Lengths of the knee joint. Other joints are in neutral position.

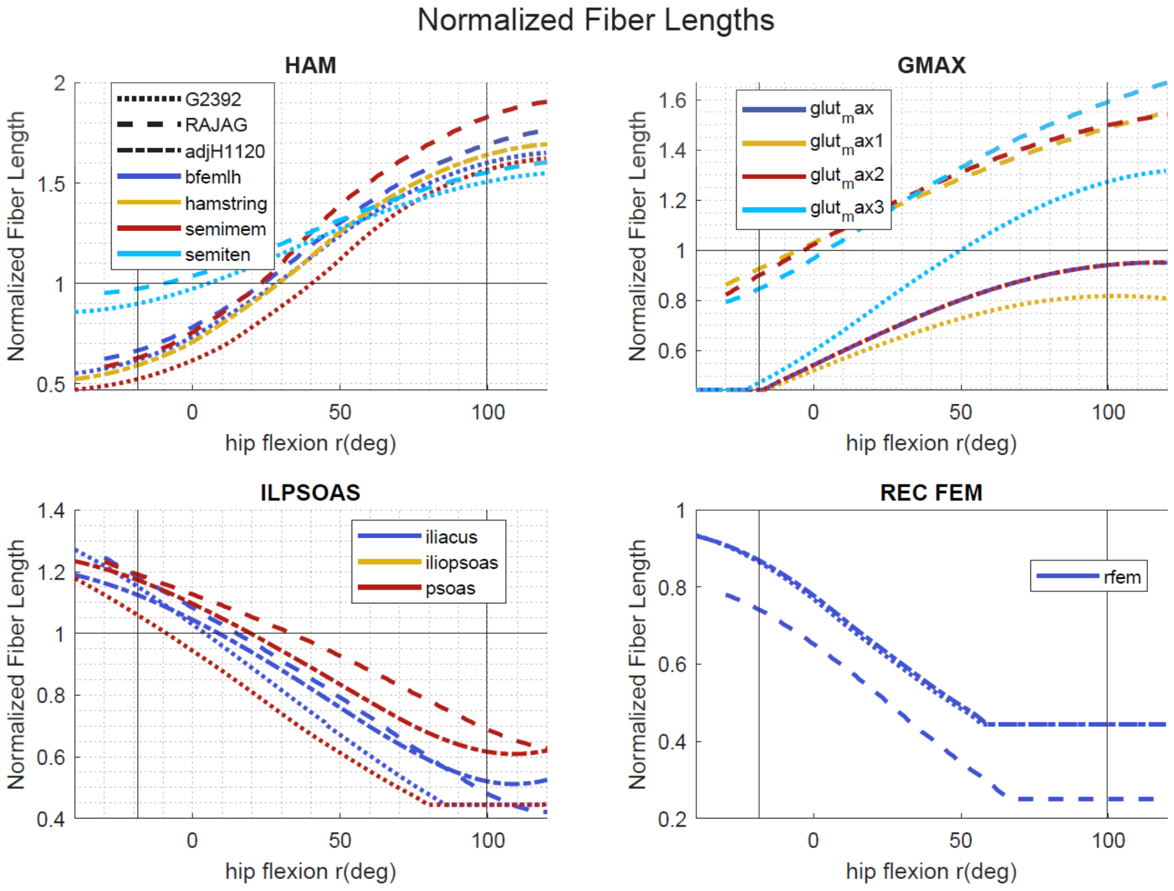


Figure 6 – Normalized Fiber Lengths of the hip joint. Other joints are in neutral position. Normalized fiber length of the Gluteus Maximus follows the G2392 shape. This means that the optimal fiber length for this muscle occurs at a much higher hip flexion (~70deg) than for the RAJAG model (0 deg) (with the knee angle at 0 degrees). The normalized fiber length of the Rectus Femoris flattens at a hip range of >~40 degrees (with the knee angle at 0 degrees). This is a lower than for the RAJAG model (~60 deg)
.

**NORMALIZED PATHWAY**

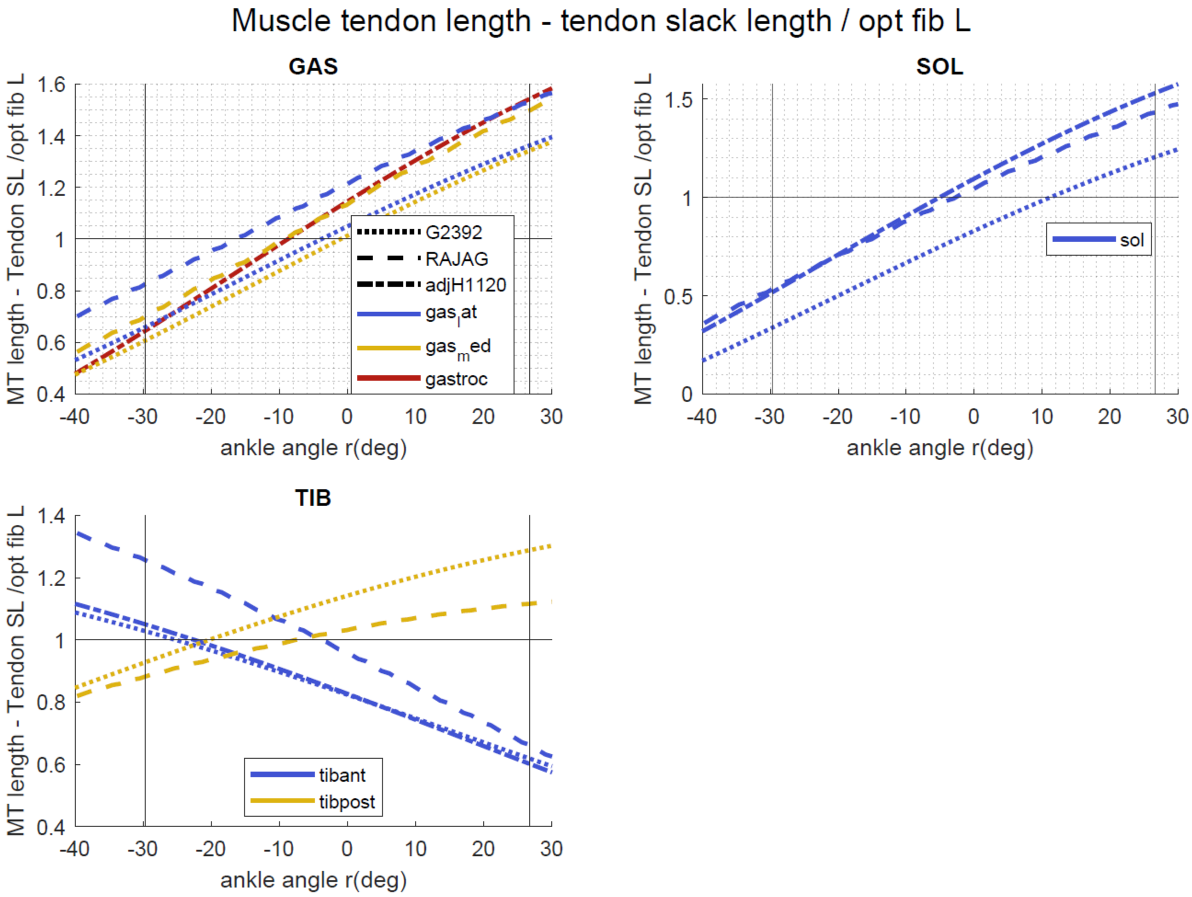


Figure 7 – Normalized Path way of muscle crossing the ankle joint. Other joints are in neutral position. The normalized pathway is estimated by subtracting the tendon slack length from the muscle tendon length and dividing by the optimal fiber length.

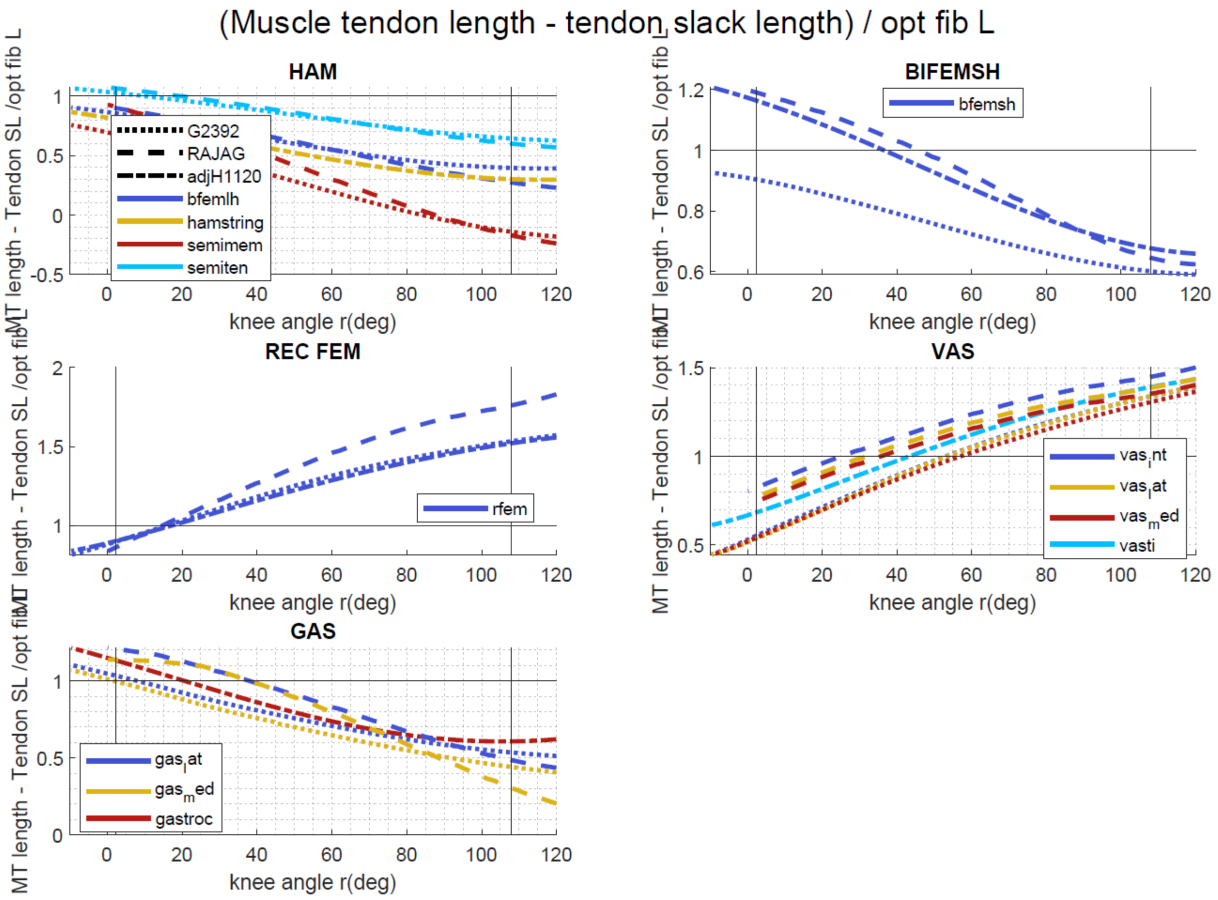


Figure 8 – Normalized Path way of muscle crossing the knee joint. Other joints are in neutral position. The normalized pathway is estimated by subtracting the tendon slack length from the muscle tendon length and dividing by the optimal fiber length. The BFEM follows the G2392 pathway which significantly differs from the RAJAG model. The RFEM follows the G2392 pathway which significantly differs from the RAJAG model.

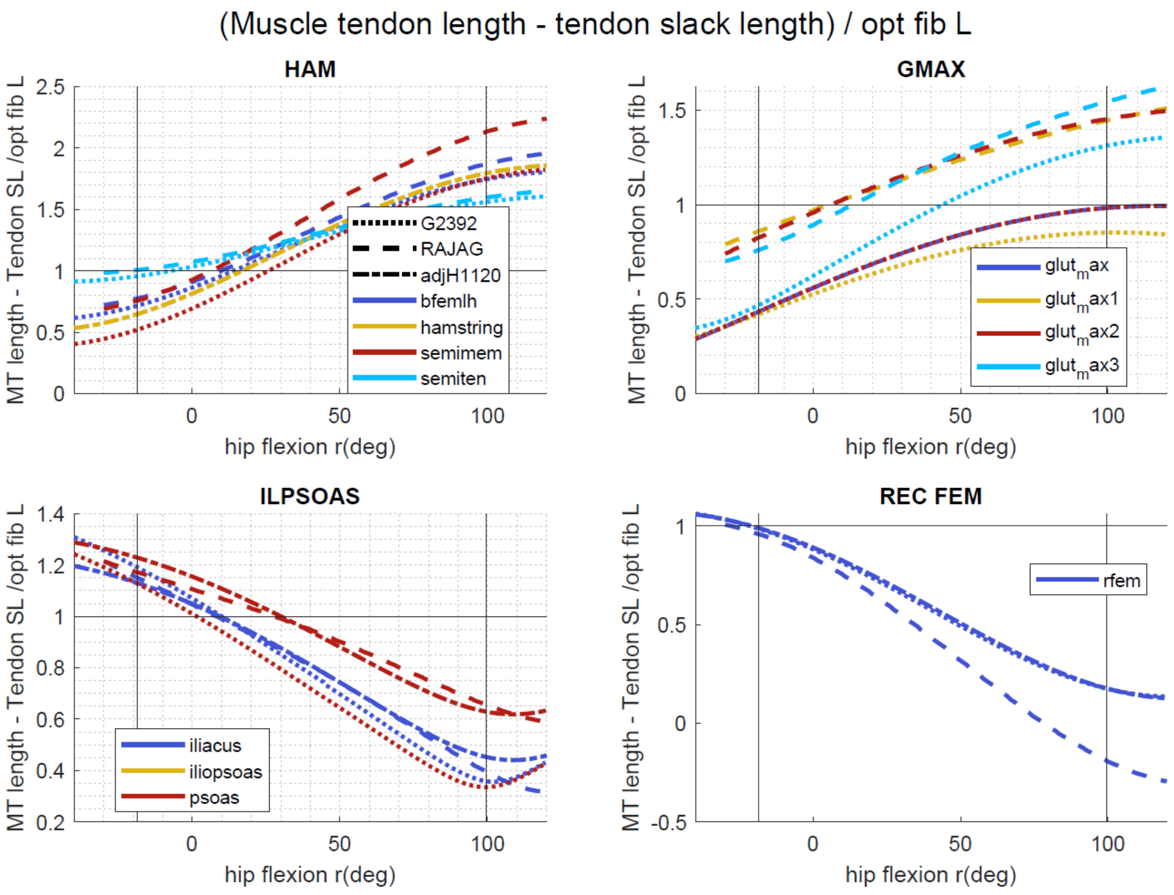


Figure 9 – Normalized Path way of muscle crossing the hip joint. Other joints are in neutral position. The normalized pathway is estimated by subtracting the tendon slack length from the muscle tendon length and dividing by the optimal fiber length.
